# Neuronal Population Encoding of Identity in Primate Prefrontal Cortex

**DOI:** 10.1101/2022.06.26.497629

**Authors:** KK Sharma, MA Diltz, T Lincoln, ER Albuquerque, LM Romanski

## Abstract

The ventrolateral prefrontal cortex (VLPFC) shows robust activation during the perception of faces and voices. However, little is known about what categorical features of social stimuli drive neural activity in this region. Since perception of identity and expression are critical social functions, we examined whether neural responses to naturalistic stimuli were driven by these two categorial features in the prefrontal cortex. We recorded single neurons in the VLPFC, while two macaques viewed short audiovisual videos of unfamiliar conspecifics making expressions of aggressive, affiliative, and neutral valence. Of the 285 neurons responsive to the audiovisual stimuli, 111 neurons had a main effect (two-way ANOVA) of identity, expression or their interaction in their stimulus related firing rates; however, decoding of expression and identity using single unit firing rates rendered poor accuracy. Interestingly, when decoding from pseudopopulations of recorded neurons, the accuracy for both expression and identity increased with population size, suggesting that the population transmitted information relevant to both variables. Principal components analysis of mean population activity across time revealed that population responses to the same identity followed similar trajectories in the response space, facilitating segregation from other identities. Our results suggest that identity is a critical feature of social stimuli that dictates the structure of population activity in the VLPFC, during the perception of vocalizations and their corresponding facial expressions. These findings enhance our understanding of social behavior beyond the temporal lobe in macaques and humans.

**Significance Statement:** Primates are unique in their ability to process and utilize complex, multisensory social information. The brain networks that support this are distributed across the temporal and frontal lobes. In this study, we characterize how social variables like identity and expression are encoded in the neural activity of the ventrolateral prefrontal cortex (VLPFC), a prefrontal region of the macaque brain. We found that single neurons do not appear to encode these variables, but populations of neurons display similar activity patterns that are primarily differentiated by the identity of the conspecific that a macaque is attending to. Furthermore, by employing dynamic, multisensory stimuli, our experiment better approximates real world conditions, making our findings more generalizable.

## Introduction

Facial and vocal expressions are a fundamental component of communication in humans and non-human primates (Darwin, 1965; Ekman & Friesen, 1971; Parr & Heintz, 2009; Partan, 2002). In macaques, viewing expressions produces robust activity through the occipital, temporal, and ventrolateral prefrontal cortices (Russ & Leopold, 2015; Shepherd & Freiwald, 2018). The ventrolateral prefrontal cortex (VLPFC) exhibits a high degree of responsivity to social stimuli, containing face cells (O Scalaidhe, 1999; O Scalaidhe et al., 1997), face patches (Tsao, Schweers, et al., 2008), neurons responsive to the angle of a viewed face (Romanski & Diehl, 2011), and neurons selective for species-specific vocalizations (Hage & Nieder, 2015; Romanski et al., 2005; Romanski & Goldman-Rakic, 2002).

While these findings suggest that the VLPFC serves a critical social function, little is known about what social variables are represented by neural activity in the region. This is in stark contrast to the temporal face patch network (Tsao, Moeller, et al., 2008), where a variety of features that drive neural responses have been identified. Anterior face patches contain neurons with selectivity for identities, whereas middle face patch activity correlates with head orientation of the viewed stimulus (Freiwald & Tsao, 2010). Patches in the fundus of the superior temporal sulcus are sensitive to changes in expression (Taubert et al., 2020) while face patches outside the traditional network display joint encoding of multiple variables, including expression and identity but also gaze and head orientation (Yang & Freiwald, 2021). Notably, information about identity is broadly distributed in the face patch network because it can be reliably decoded from small neural populations across many face patches based on representation of neurons as axes in a high-dimensional face space (Chang & Tsao, 2017). While neurons with differential responses for the angle of a viewed face have been identified in VLPFC (Romanski & Diehl, 2011), a systematic exploration of their selectivity for identity or type of expression has yet to be conducted.

A further complication in assessing the social function of the VLPFC arises from its role as a multisensory integrator. A VLPFC neuron’s multisensory response to an audiovisual face-vocalization stimulus is frequently a non-linear combination of its component responses to the face or vocalization alone (Diehl & Romanski, 2014; Sugihara et al., 2006). While this has the advantage of drastically increasing the possible set of responses a neuron or population of neurons can have (Rigotti et al., 2013), it also illustrates how response characteristics to a single sensory modality can fail to represent ethologically valid stimuli. Simply put, non-linear multisensory integration provides direct evidence that conclusions drawn from neural responses to a static image or isolated vocalization will not scale to natural versions of these inherently multisensory stimuli.

To glean more insight into the categorical social features that drive VLPFC neural activity during social communication, we recorded single unit activity in the VLPFC while macaques perceived naturalistic expressions from unfamiliar conspecifics. Given the primacy of identity and expression in similar studies (Gothard et al., 2007; Kuraoka et al., 2015; Kuraoka & Nakamura, 2007; Yang & Freiwald, 2021), we varied stimuli by conspecific identity and expression. We presented ethologically valid audiovisual movie clips from three different conspecifics, each making three discrete types of audiovisual expressions with distinct valances (aggressive, affiliative, and neutral, respectively; Hauser and Marler 1993; Partan 2002). We recorded neurons across the VLPFC and analyzed the subset that displayed consistent firing rate changes in response to at least one of the nine stimuli presented. We found that single neurons had low selectivity for single stimuli and even lower selectivity for expression or identity as a category. Rather, they responded to various stimuli over the stimulus period. Despite heterogenous firing patterns, both expression and identity could be accurately decoded from populations of neurons. Furthermore, we found that population responses to the same identity followed coherent trajectories over time that separated them from other identities in the population response space.

## Materials and Methods

### Subjects and surgical procedures

Extracellular recordings were performed in two rhesus monkeys (*Macaca mulatta*): Subject 1 (male: 12.0 kg, 7yo) and Subject 2 (male: 16.5 kg, 12yo). All procedures were in accordance with the National Institute of Health’s *Guidelines for the Care and Use of Laboratory Animals* and the University of Rochester Care and Use of Animals in Research committee. For Subject 1, prior to recordings, a titanium head post was surgically implanted to allow for head restraint and a titanium recording cylinder was placed over the VLPFC as defined anatomically by (Preuss & Goldman-Rakic, 1991) and physiologically by (Romanski & Goldman-Rakic, 2002). For Subject 2, several Floating Microarrays (FMAs, Microprobes for Life Science, Gaithersburg, MD, USA) were designed specifically for the subject’s prefrontal cortex and directly implanted into VLPFC during open surgery (inferior to the principal sulcus, anterior to the inferior limb of the arcuate sulcus).

### Experimental Design

#### Stimuli and Apparatus

In our stimulus presentation paradigm, subjects attended to naturalistic, audiovisual communication calls with their accompanying facial expressions. These vocalization movies were extracted from recordings of unfamiliar conspecifics in our home colony obtained by L.M.R. Two stimulus lists were designed such that they each contained three identities (unfamiliar conspecifics) making three ethologically distinct expressions (Figure 1B), which differed at three levels of valance: aggressive (*pant threat*), affiliative (*coo*), and neutral (*grunt*) (Hauser & Marler, 1993; Parr & Heintz, 2009; Partan, 2002). As such, our selection of stimuli renders the type of expression (e.g. *coo*) synonymous with the valance (e.g. affiliative) and expression types are henceforth referred to by their valance. Both lists contained the same identity conspecifics, who made different exemplars of the same vocalization and facial expression. Lists alternated between recording sessions, such that the population of neurons recorded contain a mix of neural responses to one list or the other; however, the abstracted category of identity and expression were preserved in the stimuli from day to day (i.e. the second list was a copy of the first with different examples of the same identity-expression paring). The movie stimuli varied in length, based on the natural length of the vocalization and associated facial movement (average length 900ms). Additional lists of visual stimuli including objects, patterns, scenes were used during preparatory phase of a recording session.

**Figure 1:**
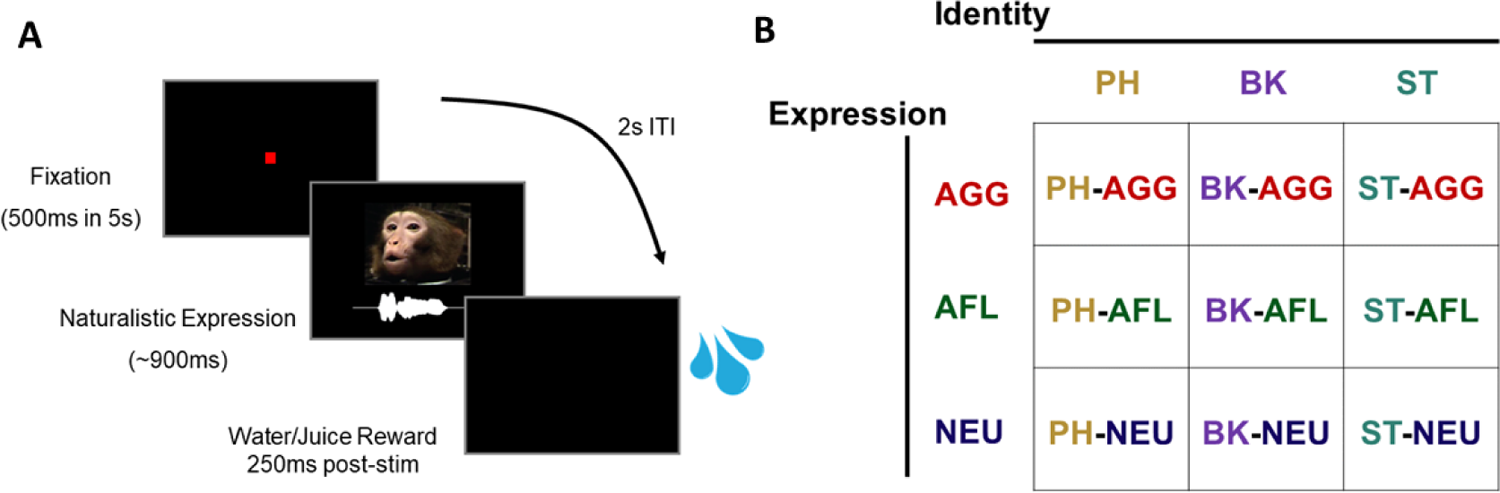
Naturalistic Expression Presentation Task 2 macaque subjects viewed movie clips (audio and video) of expressions from unknown conspecifics. A) Stimulus presentation was initiated by the subject by fixating on a central dot for 500ms. An audiovisual expression was subsequently presented (900ms mean length) and a reward was given if fixation was maintained on the stimulus through its length. (B) Matrix of stimuli showing that movie clips of 3 conspecifics (PH, BK, and ST) making aggressive (AGG: pant threat), affiliative (AFL: coo), or neutral (NEU: grunt) expressions were presented pseudo-randomly. Two lists were constructed with these parameters and alternated between recording sessions.

The audiovisual movie stimuli were processed using Adobe Premiere (Adobe Systems, San Joe, CA, USA), Jasc Animation Studio (Corel, Minneapolis, MN, USA), and several custom and shareware programs. Auditory and visual components of the movies were separated into wav and mpeg streams for processing. The visual mpeg stream was edited to remove extraneous and distracting elements in the viewing frame. The audio track was also filtered to eliminate background noise if present using MATLAB (MathWorks, Natick, MA, USA) and SIGNAL (Engineering Design, Cambridge, MA, USA). The two streams were then recombined for presentation during recordings. Stimulus audio was presented at 65-75dB SPL at the level of the subject’s ear. Movies subtended 8-10° of visual angle and were presented at eye level in the center of the computer monitor.

Neurophysiology recordings were performed in a sound-attenuated room lined with Sonex (Acoustical Solutions, Richmond, VA, USA). Stimuli were presented through a video monitor flanked by speakers with ±3 dB, 75-20,000 Hz frequency response (PH5-vs, Audix, Wilsonville, OR, USA). The monitor and flanking speakers were located 29 inches from the subject’s head and centered at the level of the subject’s eyes. During all neural recordings, eye position was monitored using an infrared pupil monitoring system (ISCAN, Burlington, MA, USA).

### Experimental Procedures and Presentation Task

Animals were acclimated to the laboratory and testing conditions. Subjects were then trained on a version of the audiovisual stimulus presentation task that did not include the stimuli used for recording (though they did encounter some stimuli in previously trained/recorded tasks). For Subject 1, the head was restrained by means of the surgically implanted post and a custom-built fixation system mounted to the primate chair. Then, either a single recording electrode (tungsten, FHC Inc., Bowdoin, ME, USA) or linear array (16ch or 32ch, V-Probe, Plexon Inc., Dallas, TX, USA) was lowered into the VLPFC each day, using a hydraulic microdrive and XY positioner (MO-95C, Narishige Int. USA, Amityville, NY, USA) that mounted onto the recording cylinder. For Subject 2, the head was temporarily restrained with a moldable plastic headpiece while the recording headstage was connected to the FMA Omnetics connectors. Once connected, the plastic headpiece was removed and the subject was free to move through a limited range in the chair during recordings.

During the preparatory phase of the recordings and cylinder preparation, subjects viewed lists of visual objects, patterns, and scenes. For Subject 1, an electrode or linear array was lowered into the VLPFC while the subject viewed these preliminary lists. The linear array was lowered until all of the contacts were below the dural surface. Single electrodes were lowered past layer 1 until single cells could be easily isolated. For Subject 2, the implanted arrays were surgically implanted so no manual modifications could be made and 64 channels of data were recorded by default.

Wideband neural data and behavioral event codes were recorded, digitized at 40,000hz, and saved to disk using a neural data acquisition system that featured analog to digital conversion at the head stage (Omniplex, Plexon Inc.). For Subject 1, the wideband signal recorded from Plexon v-probes or Tungsten electrodes was bandpass filtered to 600-6000hz and played back through a pre-amplifier and speaker in order to optimize the position of the electrode, prior to recording. In certain instances, the 600-6000hz “spikeband” was also saved alongside the wideband data.

At the start of each trial, a red fixation point (0.15×0.15 degrees) was presented at the center of the monitor. Subjects were given a 5s period in which a 500ms sustained fixation on the point would trigger the beginning of the task and a vocalization movie was presented. During the movie, subjects had to maintain their gaze within the viewing window corresponding to the boundaries of the stimulus (10°x10° window) for the entire duration of the movie stimulus. Eye position was monitored continuously through the trial. Successful completion of the trial resulted in a juice or water reward 250ms after the end of the stimulus (Figure 1). Failure to establish 500ms of fixation within the given time or failure to maintain gaze within the boundaries of the stimulus was considered a failed trial and immediately terminated the presentation of the stimulus. All successful trials were equally rewarded and unsuccessful trials were stopped upon loss of fixation. Trials were presented pseudo-randomly for 10-12 repetitions (Subject 1) or 15 repetitions (Subject 2) of each stimulus. The inter-trial interval (ITI) was 2 seconds. The timing of behavioral contingencies, presentation of stimuli, delivery of reward, and monitoring of eye position were controlled by custom software running on a dedicated computer.

### Data Analysis

#### Spike Sorting

Spike sorting was conducted manually, offline, after recording sessions were completed using a dedicated computer and software (Offline Sorter, Plexon Inc.). Spikes were either sorted from the wideband signal that was high-pass filtered at 600hz (4-pole Butterworth) or from saved spikeband data (see Presentation Task and Experimental Procedures). The generalized process for spike sorting started with thresholding the filtered wideband or spike band at a level where clusters of spikes were visually distinct in Principal component (PC) 1/2, 2/3, or 1/3 spaces. Spikes were then realigned to their global minimum after the first threshold crossing. Clusters were preliminarily designated as a single unit within one of the PC spaces and then corroborated through clustering in another PC space. In rare instances, a cluster that could not be corroborated in another PC space would be found in a PC x Non-linear energy or PC x Peak-Valley space. Uncorroborated clusters had their unit designation removed and were thus not identified as single units in our analysis. Clusters that had greater than 0.2% inter-spike interval (1ms) violations were reduced in size, typically in the corroborating space, to meet the threshold. If such a threshold could not be met, the unit designation was removed. As such, the single units analyzed in this study have, at most, 0.2% ISI and can be visually identified as clusters in at least two combinations of PC 1, 2, 3, Non-linear energy, or Peak-Valley spaces.

#### Identification of Responsive Units

Because our stimuli varied in duration (range 400 – 1600, average 900) due to the natural length of the associated vocalization, any period of time used to identify responsive units represented a trade-off between including time after a stimulus had terminated (for shorter stimuli) or cutting off a response before a stimulus had terminated (for longer stimuli). We found units that had stimulus-dependent response latencies (Figure 2). Previous VLPFC studies indicate that response latencies may vary between 50 – 450 msec. Therefore, we decided to look for responses over an inclusive time period and assessed responsivity to stimuli between 0-1500ms, after stimulus onset. This period of time is henceforth referred to as the “stimulus period”. While this was 100ms shorter than our longest stimulus, the majority of cells responsive at late time points were also responsive at earlier ones for shorter stimuli, which facilitated their identification using the analyses described below.

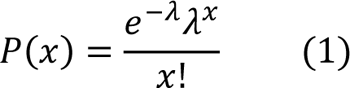

**Figure 2:**
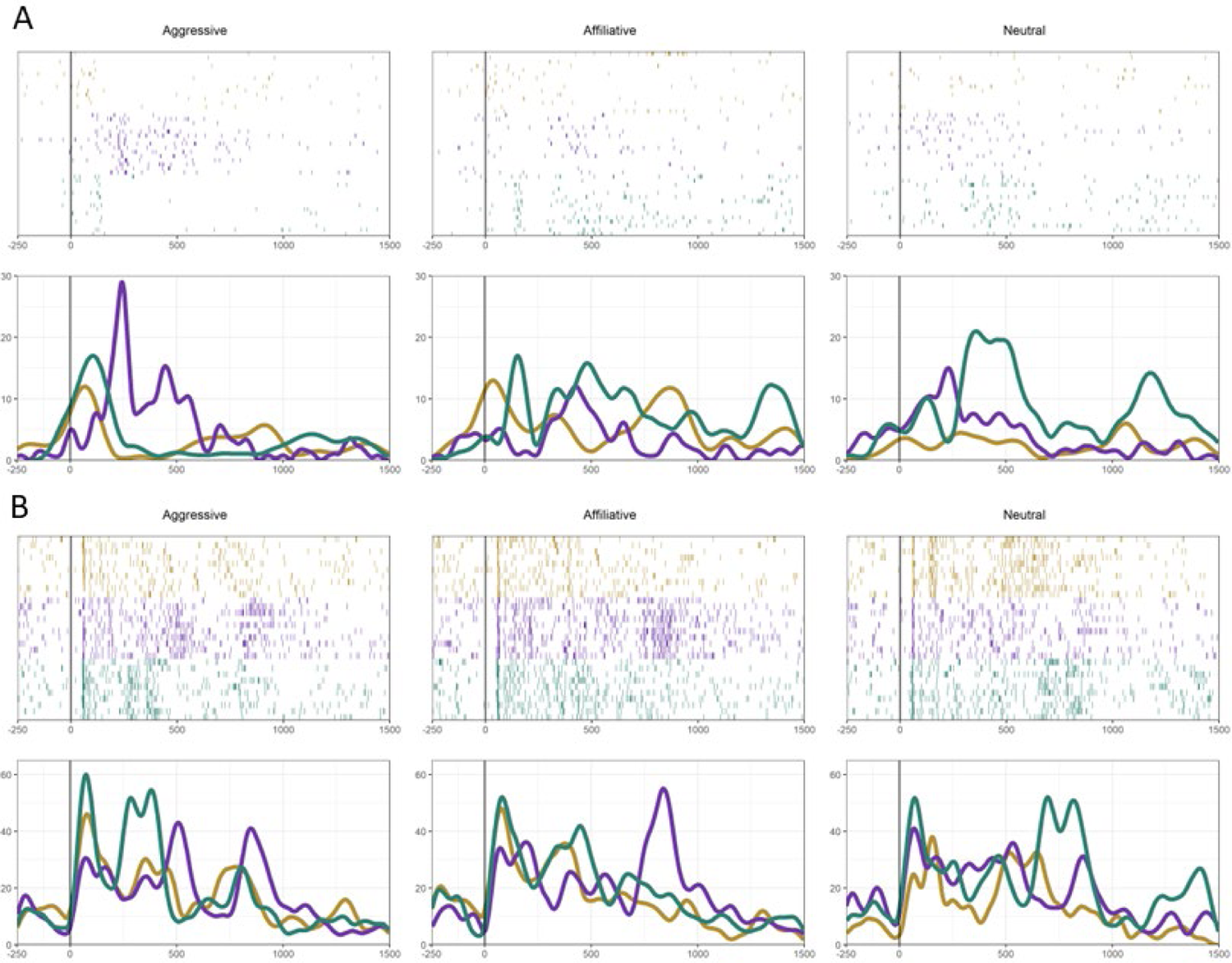
Single Units Exhibit Varied Responses Representative responses to multiple repetitions of all nine expressions are grouped by expression. Responses to identity PH are in gold, BK in purple, and ST in teal. Raster plots for each trial are presented above with corresponding SDFs below, grouped by identity. 0ms represents the onset time of the stimuli. Because these stimuli were naturalistic expressions, consisting of a video and audio component, stimuli varied from 400ms to 1000ms, based on the duration of the expression. A.) Identity was a significant modulator of the firing rate for this unit, as well as the interaction between Identity and Expression (Two-Way ANOVA Identity X Expression, p < 0.005 for Identity, p = 0.154 for Expression, p < 0.005 for Interaction). B.) Another responsive unit. In this case, Identity, Expression, and their interaction significantly modulated the firing rate (Two-Way ANOVA Identity X Expression, p<0.005 for Identity, p=0.025 for Expression, p<0.005 for Interaction).

Responsive units were identified using a sliding bin analysis to determine time bins where the Poisson probability (Equation 1) of the average spike count for a given condition was higher than the mean baseline spike count. For each unit, a mean spike count for the 200ms prior to every stimulus presentation was calculated (λ). Then a sliding bin analysis was conducted in 200ms time bins from 0-1500ms post-stimulus onset, shifting by 100ms, to derive the mean spike count in each 200ms bin for each stimulus presentation (10-15 repetitions, spike count rounded to the nearest whole number). Equation 1 was used to calculate the Poisson probability of spiking within each bin, where λ was the mean spike count in the 200ms prior to stimulus onset across all stimuli, *x* was the mean spike count in a particular 200ms bin for a single stimulus of interest, and *P*(*x*) was the probability of that the spike count *x* was would be seen within the condition and bin being analyzed given the average spontaneous spike rate λ. This calculation was repeated for all bins and for each stimulus. Units with at least one bin in one stimulus condition where the probability of spike count was less than .1 were deemed responsive in this task. This analysis was conducted using the *R* programming language.

### Selectivity Analysis

For each of the 285 responsive units, a selectivity index (*SI*) was calculated with regard to the selectivity for specific stimuli within our stimulus set, the identities in the set, and the expressions in the set. The index provides a 0-1 scale for each unit, through which the selectivity for a particular stimulus or category of stimuli can be quantified. In this analysis, a unit that selectively fired for a single audiovisual stimulus, expression, or identity would have an SSSS = 1 and a unit that fired uniformly to all instances would have an *SI* = 0. Equation 2 shows the calculation:

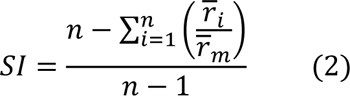

where *r_i_* represents the mean of the absolute value of the difference in firing rate between the baseline (−200ms - 0ms) and response period (0 - 1500ms) for a given stimulus, identity, or expression type i; r_m_ is the maximum of r_i_ for all i; and *n* is the number of stimuli (9), identities (3), or expressions types (3) for which an *SI* was calculated.

### Two-Way ANOVA for Expression and Identity

To assess the influence of expression type and identity on the firing rates of responsive units, we fit the firing rate of each unit during the response period (0-1500ms) with a Two-Way ANOVA model of the form in Equation 3:

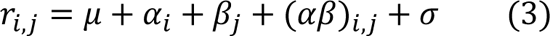

where *r*_i,j_ is the firing rate over all expression types (i = 1,2,3) and identity (j = 1,2,3) levels, μ is the mean firing rate, α is the effect due to the type of expression, β is the effect due to identity, (αβ) is the effect due to the interaction of expression type and identity, and σ is Gaussian noise. This ANOVA was performed for each responsive unit, rendering p values for the statistical significance of the effect of expression type (α), identity (β), and interaction (αβ) on each unit’s firing rate. Units were thresholded as positive for each effect at the p=0.05 level for that effect. As such, units could be significant for multiple effects. Two-Way ANOVA analysis and plotting was completed using *R* and packages *stats*, *ggplot2*, and *ggVennDiagram*.

### Single Unit K Nearest Neighbor Decoding

For all 285 responsive units, a *k* nearest neighbors classifier (KNN) was used to decode the expression type or identity of a stimulus in a single trial based on the firing rate observed in that trial. For each trial that was classified, the Euclidean distance between the firing rate in the trial to be classified and all trials in a training set were calculated. The trial was then classified based on the majority classification in an odd number of nearest neighbors within the training set. Classifications were made and accuracy was calculated using 5-21 nearest neighbors. The maximum accuracy across this set of nearest neighbors was used as the accuracy for the unit being analyzed. This analysis was conducted with 3-fold cross validation (CV) with 5 repetitions within each fold averaged to produce that fold’s accuracy metrics. The matrix provided to the decoder was *n* rows by 1 column + a column of labels, with *n* = number trials (typically 90 or 135 for 10 or 15 repetitions of each stimulus, respectively) and the column containing the 0-1500ms firing rate for a single unit. This analysis was implemented in *R* using additional packages *caret*, *knn*, and *tidyverse*.

### Pseudo-population K Nearest Neighbor Decoding

Pseudo-populations of sizes 2-280 neurons were constructed from the responsive units. Expression type and identity were then decoded from the pseudo-population using a KNN model, similar to that used for single unit decoding. In this case, classifications were made and accuracy was calculated using 5, 7, or 9 nearest neighbors and 2-fold CV. 50 repetitions were conducted at each pseudo-population level and each repetition included a random selection of units to include in the population and randomization of trials within the pseudo-population, in addition to standard randomization of the data for CV.

For each pseudo-population size, a matrix of 135 rows by *u* columns was created, containing the z-scored firing rates of units to 135 trials as rows (15 repetitions of 9 stimuli). In cases where 135 trials were not recorded for the unit, additional data was produced by random sampling from existing data for that unit. In the 5 cases where there were an unbalanced number of completed trials for a unit (e.g. a unit had 10 repetitions of one stimulus and 9 of another) the unit was removed from this analysis. The order of trials was shuffled prior to insertion in the matrix. *u* was the pseudo-population size and units from the 280 responsive units appropriate for this analysis were sampled at random to populate *u* columns on each iteration of the decoding analysis.

In summary, a 135 x *u* data matrix + a column of labels was the input data for the KNN decoder. Euclidean distance was used, 2-fold CV was completed on each iteration, and 50 iterations were completed for each pseudo-population size *u*. The data matrix was rebuilt, starting with random selection of units, for each iteration. This analysis was implemented in *R* using the previously mentioned packages.

### Decoding of Identity and Expression Over Time

A pseudo-population of 285 was created from all responsive units by creating a 135 x 285 data matrix + a column of labels (expression type or identity) for 200ms bins (200ms bin size, 100ms step size, 16 total bins) across the response period. A single column represented the z-scored firing rate in a 200ms period of interest for a single unit in response to each of 135 trials. Random sampling of existing data was used to produce 135 trial column vectors for units that did not have 135 recorded trials (same as previous). The order of trials was shuffled for each unit prior to insertion into the matrix (same as previous). Decoding was performed on firing rates in 200ms time bins, sliding forward by 100ms, starting at 250ms before stimulus onset and ending with a bin starting at 1350ms after stimulus onset.

For each 200ms bin, the decoder generated mean vectors for each classification (either expression type or identity) from the training data and classified each trial in the test data based on the mean vector it is maximally correlated with (Pearson’s correlation coefficient). This was done using 2-fold CV and repeated for 50 iterations for each 200ms bin. Mean accuracy and standard error were derived from the 50 iterations. This analysis was conducted in *MATLAB* using the *Neural Decoding Toolbox* (Meyers, 2013) and visualized in *R* using packages previously mentioned.

### Principal Component, Group Distance, and Trajectory Analysis

The mean firing rate of each of the 285 responsive units was calculated for each of the 9 stimuli using the 0-1500ms response window. The rates were then z-scored and compiled in a matrix of 9 rows x 285 columns (stimuli by units). Principal component analysis (PCA) was conducted on this matrix to project the population response to each of these 9 stimuli into 8 retained dimensions. When visualized, the coordinates of the 9 population responses were taken from their respective PC.

Distances between population responses to stimuli were calculated using the Euclidean distance between the coordinates in the preserved number of PCs. For example, the distances in a population space preserving 2 PCs were calculated for each point using 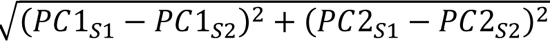 where *PC*1_S1_ is the coordinate along PC1 of the population response to stimulus 1. These distances were grouped according to whether they were a distance between the same identity (ID), between the same expression type (EXP), or between mismatches of identity and expression (OTHER). A Two-Way ANOVA of the form in Equation 4 was fitted to the distances calculated in spaces retaining 1-8 PCs. A post-hoc test, Tukey’s Honest Square Differences (Tukey HSD) was used to specify which groups had statistically significant differences from each other.

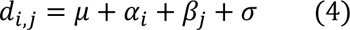

The trajectory of the neural population response to each stimulus was visualized by projecting the position of the population response at each time point of interest into the 8 dimension PCA space of the population at the final time point of 1500ms. First, the rotation matrix mapping the original space to the 8 PC space (**R**) was calculated from the PCA covering the full time range of 1500ms. For each time point of interest (ii, 50ms increments from 0m-1500ms), a new population matrix (**P**_i_) of 9 rows x 285 columns (stimuli X units) was produced from the mean firing rate to each stimulus from 0 - i ms post stimulus onset. This matrix was rotated into the final PC space (**P**_i_**R**) and visualized by retaining the respective PCs. This was repeated for all *i* intervals of interest to produce visualization of the path the population response to a single stimulus took over time. These analyses were conducted using *R* and additional packages *PCA*, *factominer*, and *factorextra*.

## Results

### Single Units Responsive to Naturalistic Stimuli Exhibit Complex and Non-Selective Responses

We recorded and isolated 466 units from the VLPFC of 2 macaques, while they viewed audiovisual movie clips of 3 different conspecifics making 3 different types of vocalization-expressions: either an affiliative, aggressive, or neutral expression. In the subsequent analysis, “expression” and “expression type” refers to the type of vocalization-expression (*coo*/affiliative, *pant threat*/aggressive, or *grunt*/neutral) whereas “stimulus” or “stimuli” refers to the specific identity-expression combinations i.e. one or multiple of the 9 audiovisual movie clips seen and heard. Due to variability of responses to stimuli found across time (Figure 2), a custom filter for responsivity, using a sliding bin analysis of Poisson probability, was constructed to identify units responsive in this task (see Methods: Identification of Responsive Units). From this analysis, all units identified as responsive had at least one 200ms period where the firing rate was significantly elevated or reduced compared baseline, in response to at least one stimulus. Additionally, units that had fewer than 10 repetitions of all stimuli were removed from this analysis.

285 units met the aforementioned criteria (n=172 subject 1, n=113 subject 2). Figure 2 depicts the typical activity patterns found across the population of responsive units. Units had peak responses that varied in time or had multiple peaks across the response period and displayed varying activity patterns relative to the specific stimulus presented.

The observed dynamic firing patterns prompted a quantitative understanding of the selectivity of single units for specific stimuli, identities, and expression types. A Selectivity Index (SI) was calculated for the firing rate across 0-1500ms for each unit, with regards to selectivity for each stimulus, each identity, and each expression (see Methods: Selectivity Analysis). Using this index, each neuron’s response to a condition can be expressed along a 0-1 scale, with 1 signifying exclusive responsivity to a single condition and 0 signifying uniform responsivity to each condition. Neurons responsive to our task were only moderately selective for specific stimuli (mean SI: 0.41 ± 0.13) and less selective for identity (0.22 ± 0.13) or expression type (0.23 ± 0.13) (Figure 3). The increased selectivity for stimuli compared to identity or expression was statistically significant (Kruskal-Wallis followed by pairwise Wilcoxon rank sum, p < 0.001, Bonferroni corrected). This suggests that neural firing of single units in response to naturalistic expressions correlates more with combinations of stimulus features than categorical variables like identity or expression type.

**Figure 3:**
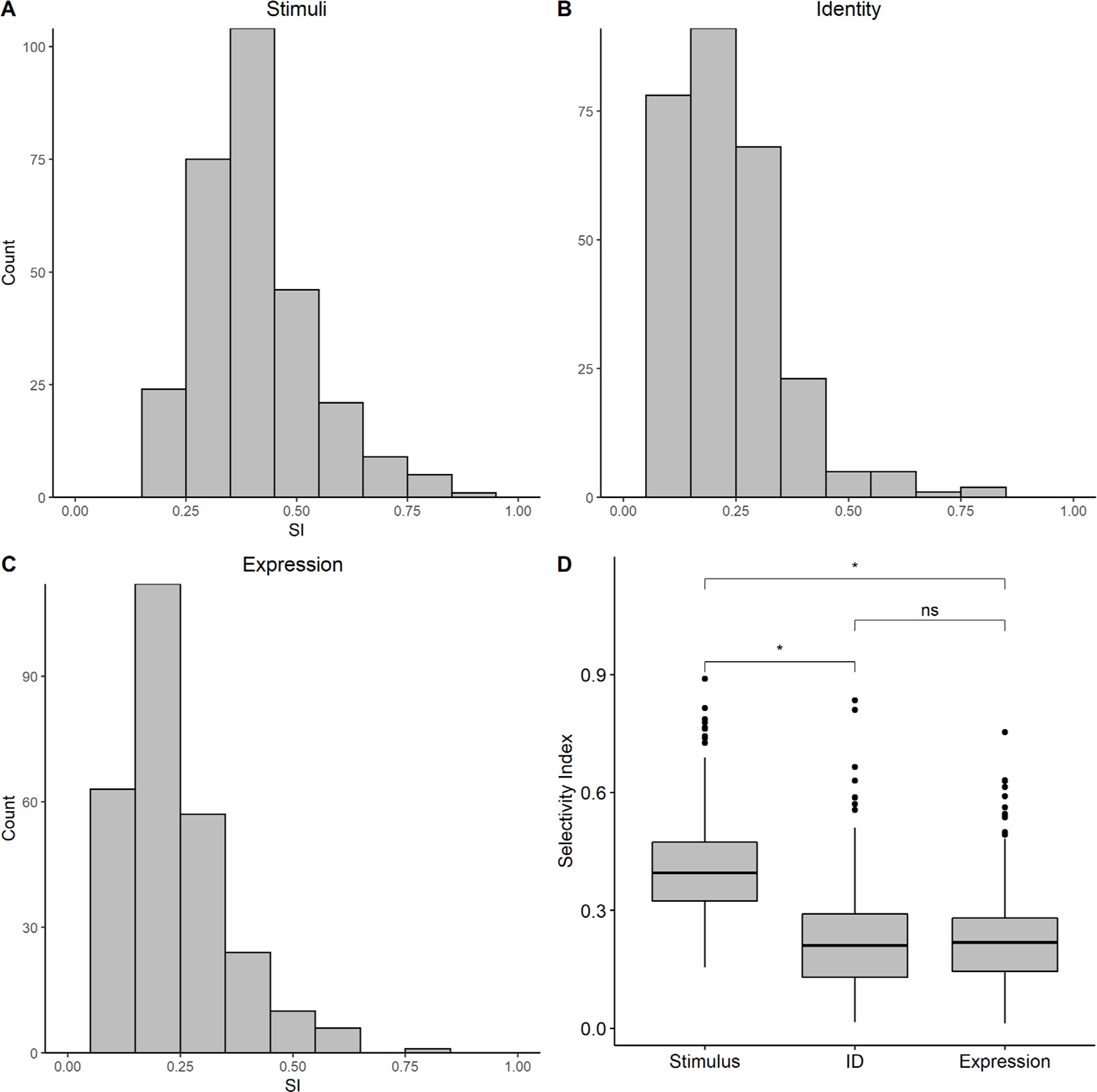
Selectivity Index Distributions A selectivity index was calculated for all 285 units with regards to selectivity for one of nine stimuli, one of three identities, or one of three expressions. The plots above depict the counts of cells at each tier of selectivity index (in bins of size .1), with regard to selectivity for stimuli (A), identity (B), and expression (C). The mean of each distribution was significantly different from 0 (3 Wilcox tests, p < 0.005 for Stimuli, Identity, and Expression means compared to 0). Boxplots in (D) depict a comparison between means of SI for each factor of interest. The population was more selective for specific stimuli (0.41 ± 0.13) compared to identity (0.22 ± 0.13) and expression (0.23 ± 0.13) (Kruskal-Wallis, df = 2, p < 0.001 followed by pairwise Wilcoxon rank sum, p < 0.001, Bonferroni corrected). While these units are not uniformly responsive, they are also not highly selective for specific stimuli, identities, or expressions.

Despite having a higher *SI* for stimuli, the selectivity for expression type and identity across the population of units was not 0; therefore, units did not respond uniformly or indiscriminately to all identities or expressions, but they were also not selective for either variable. Thus, it was possible that these categorical variables influenced firing rates in a more subtle way. In order to further characterize the potential influence of these factors on single unit firing rates, we conducted a Two-Way ANOVA analysis modeling expression type, identity, and the interaction of both factors as predictors of firing rate in the 0-1500ms stimulus period. A Two-Way ANOVA was conducted for each unit, and we found that 111 units from our population had a significant effect of expression type, identity, or their interaction (Figure 4A). Within the set of 111 units, 80 units had a significant effect of identity, 56 had a significant effect of expression, and 66 had a significant effect of their interaction (Figure 4B, p < 0.05). It should be noted that these counts represent overlapping subsets, because a single unit could have multiple significant factors in the Two-Way ANOVA; this is portrayed in the Venn Diagram in Figure 4C. The classification with the highest number of units was those for which there was a significant effect of identity, expression, and their interaction (30 units, 27% of the 111 significant units). The next highest group was those with a significant effect of identity only (28, 25%). Firing rates and *SI* for two units that showed ANOVA effects of expression type, conspecific identity, and interaction are shown in Figure 5. Overall, these results provided evidence that expression type and conspecific identity influenced the firing rates of a subset of single units.

**Figure 4:**
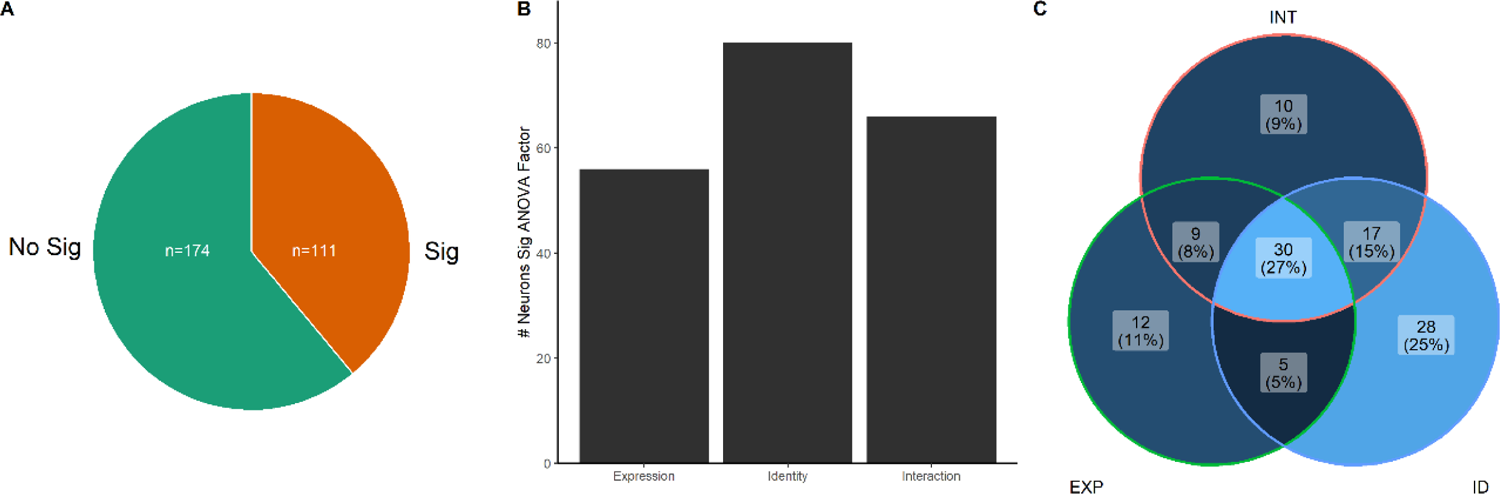
Two-Way ANOVA of Single Cells Firing Rates Two-Way ANOVA of expression type (EXP), identity (ID), or the interaction (INT) of both. (A) Of the 285 responsive units, 111 were statistically significant (p<0.05) for at least one ANOVA factor (in orange), indicating an influence of EXP, ID, or INT. (B) Shows the counts of units with significant effects for each factor or interaction in this analysis. 80 units were significant for ID, 56 for EXP, and 66 for INT. (C) The units from (B) represent an overlapping population that can have multiple significant factors. Thus, there are 7 possible classifications for each unit and the distribution of these classifications is shown in the Venn Diagram. Lighter shades indicate higher counts. Percentages represent a proportion of the 111 Two-Way ANOVA positive units in (A). Overall, a co-occurrence of significant effects of ID, EXP, and INT was the highest portion in the Two-Way ANOVA positive population, indicating influences of both ID and EXP as well as specific stimuli. Examples such units with these effects are shown in **Figure 5**.

**Figure 5:**
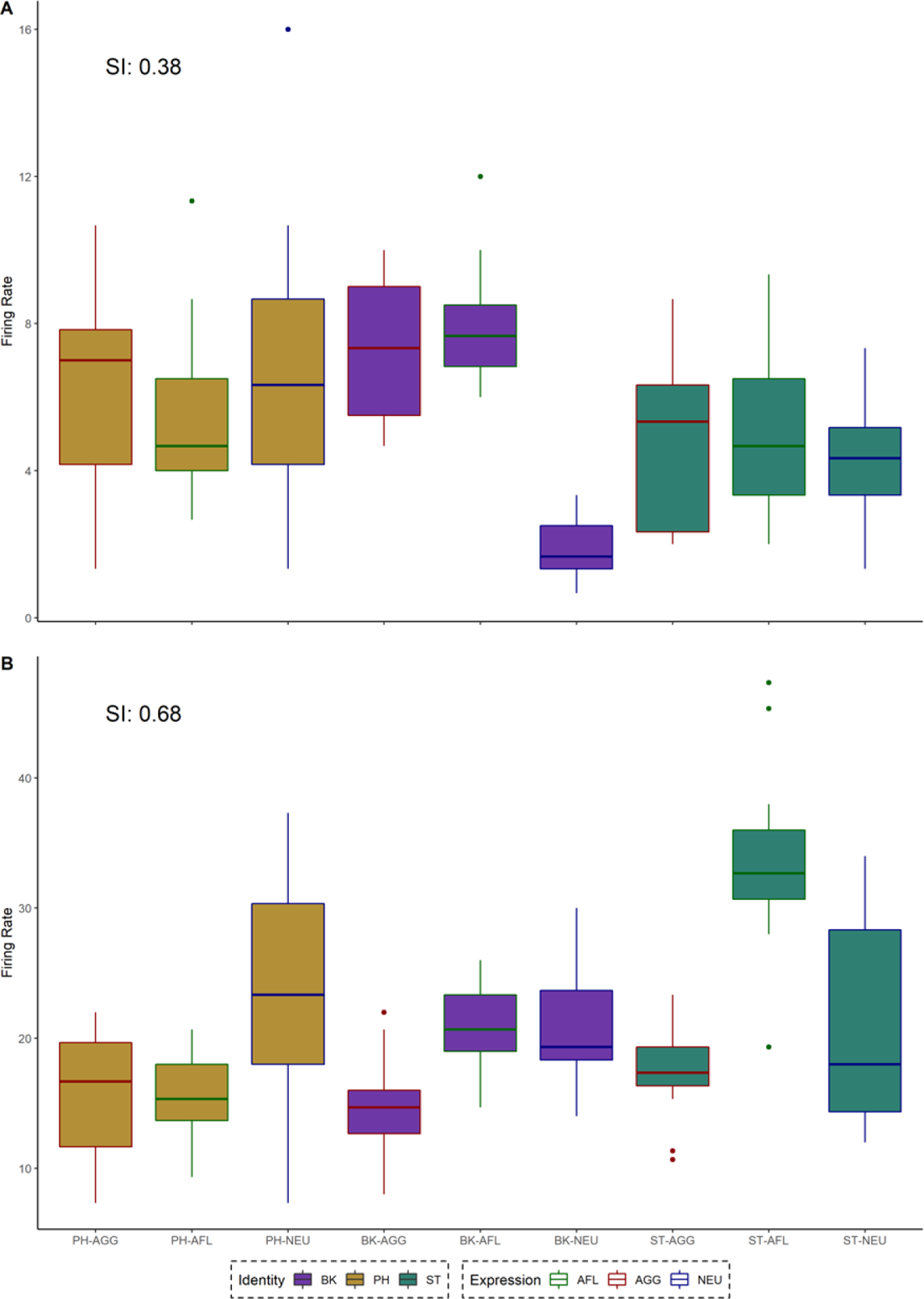
Firing Rate by Stimulus for Two Units with Effects of ID, EXP, and INT Two units with box and whisker representations of firing rates for each stimulus (horizonal bar = median, box = interquartile range (IQR), stems = 1.5*IQR). Boxplot line boarders correspond to expression and boxplot fill colors correspond to identity. In both A and B, both units had significant effects of expression, identity, and interaction in Two-Way ANOVA analysis. A Selectivity Index (SI) calculated by stimulus is provided and y axis is scaled to a relevant range for each unit. In A, a unit with firing rate trends for expressions and identity also has a single stimulus (BK-NEU) for which there is a markedly different firing rate distribution than all others. B depicts a similar trend as in A for a unit with a higher SI. In this case, stimulus ST-AFL had a markedly higher firing rate than others.

### Accurate Decoding of Expression and Identity Requires a Neural Population

Since single units paradoxically exhibited both generally low selectivity and main effects of expression type and identity, we assessed whether the effects of these variables were sufficient to differentiate expressions or identities based on the firing rates of single units. We used a decoding approach in which single trials were classified by conspecific identity or expression type based on their Euclidean distance to *k* nearest neighbors (KNN model, 3-fold cross-validation (CV) with 5 repeats, maximum decoding accuracy across k = odd integers from 5-21, see Methods: Single Unit *K* Nearest Neighbor Decoding). Figure 6 depicts the results for this decoding model. The mean expression decoding accuracy averaged across the set of 285 responsive units was 0.37 ± 0.05 (significantly above chance of 0.33, p < 0.005, Wilcoxon Rank Sum, n = 285). The comparable mean decoding accuracy for identity was 0.38 ± 0.06 (p < 0.005, Wilcoxon Rank Sum, n = 285), which was also significantly higher than the mean for expression (difference of 0.01, p < 0.005, Wilcoxon Rank Sum). These results suggest that, on average, the firing rates of units responsive to naturalistic stimuli in the VLPFC carry some reliable information regarding expression type and conspecific identity. However, though statistically significant, the average decoding accuracy for either of these variables was only marginally above chance suggesting that single unit firing rates are not a reliable source of information.

**Figure 6:**
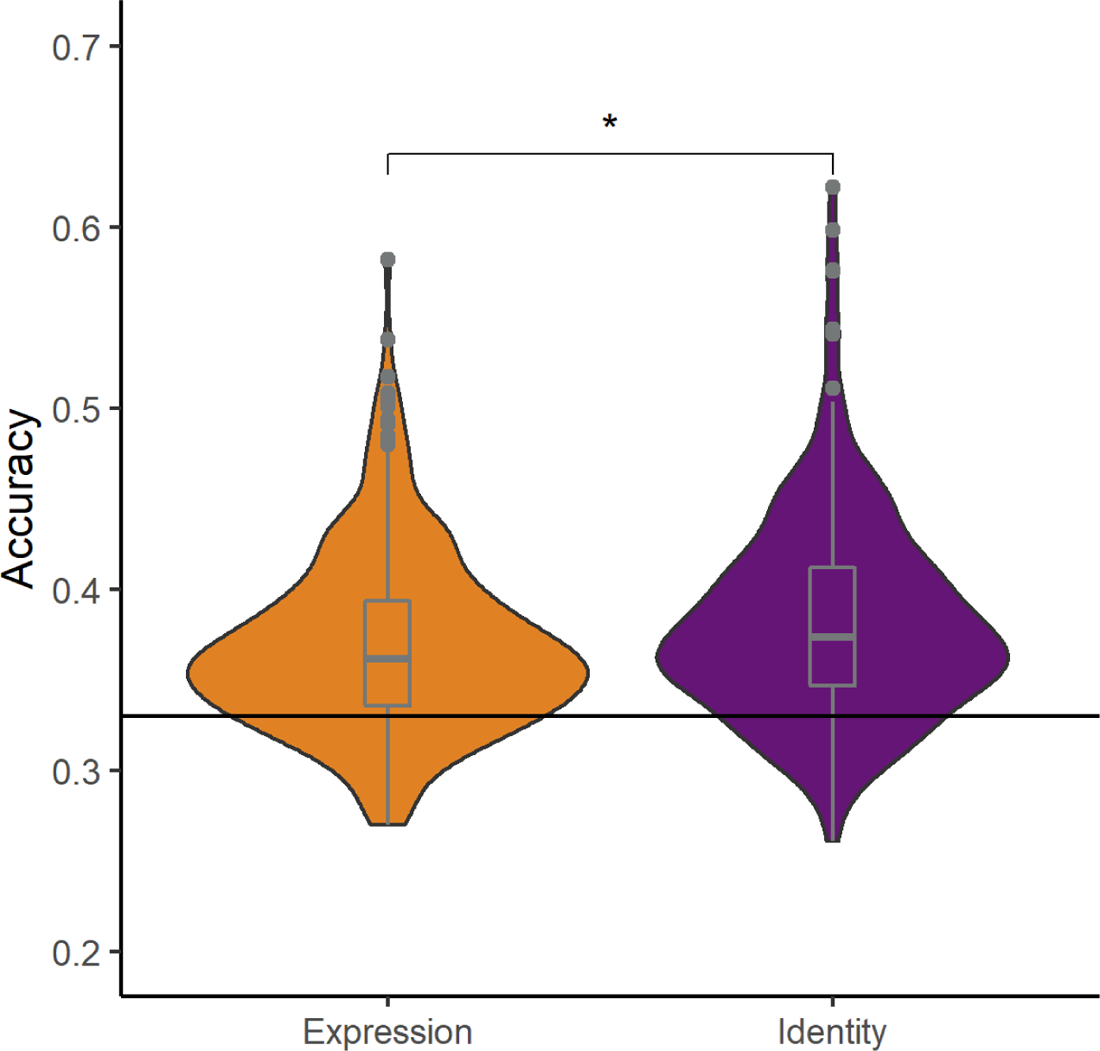
Weak Decoding of Identity and Expression from Single Unit Firing Rates For each single unit, we classified single trials based on the firing rate similarity (Euclidean distance) to 5-21 nearest neighbors in training sets (KNN model, 3-fold CV, 5 repeats). The maximum decoding accuracy across 5-21 nearest neighbors was used as the single units decoding accuracy. The mean decoding accuracy of expression (0.37 ± 0.05 SD) and identity (0.38 ± 0.06) were significantly above the chance value of 0.33 (p < 0.005). Additionally, the mean decoding accuracy for identity was above expression (difference of 0.01, p < 0.005, Wilcoxon Rank Sum, Bonferroni corrected).

Although decoding of categorical social variables using single unit firing rates was largely inaccurate, the diversity of responses found in these units may facilitate accurate decoding at the population level. To understand whether decoding of expression type or conspecific identity could be achieved by utilizing the populations of units, we conducted a decoding analysis on increasing sizes of pseudo-populations of responsive units (280 units used, 5 removed due to unbalanced number of trials). Briefly, a pseudo-population was constructed by randomly combining units to meet a target population size and randomly matching trials of the same stimuli within that population. In cases where a unit had fewer than 15 repetitions per stimulus, firing rates for that the remaining repetitions were randomly sampled from the existing data for that stimulus until 15 repetitions of each stimulus were represented in the pseudo-population for that unit. Decoding accuracy for a pseudo-population was derived from the maximum accuracy of a KNN model (see Methods: Pseudo-population *K* Nearest Neighbor Decoding) across all *k* (*k* = 3,5,7, 2-Fold CV, 50 repetitions with randomization of units in the population at each repetition). It should be noted that decoding for expression type or conspecific identity in this analysis is made more difficult by two factors inherent to the methodology of our experiment. First, single units were recorded during viewing of one of two possible sets of stimuli that contained slightly different exemplars of the same identity-expression pairings as stimuli (i.e. different exemplars of the same identity making the same expression). Thus, the decoding for conspecific identity or expression type at the population level requires a generalization of that variable across two exemplars of each stimulus. Secondly, KNN classifiers suffer from “the curse of dimensionality” (Weber et al., 1998), in which higher dimensional spaces naturally lead to sparse representations of randomness, particularly beyond 10 dimensions.

Despite these factors, the decoding accuracy for both expression and identity increased as the population size increased (Figure 7). Expression decoding increased from .40 at a pseudo-population size of 2 to .68 at 280. Identity increased from .41 at size 2 to .80 at 280. Inspection of Figure 8 shows a non-monotonic but consistent increase in decoding accuracy as a function of population size. Additionally, the accuracy of identity was consistently above that of expression for all population sizes. This evidence supports the assertion that the decoding of expression and identity requires population level activity from single units in the VLPFC. Additionally, it suggests that information about the identity of a conspecific is abundant at the population level.

**Figure 7:**
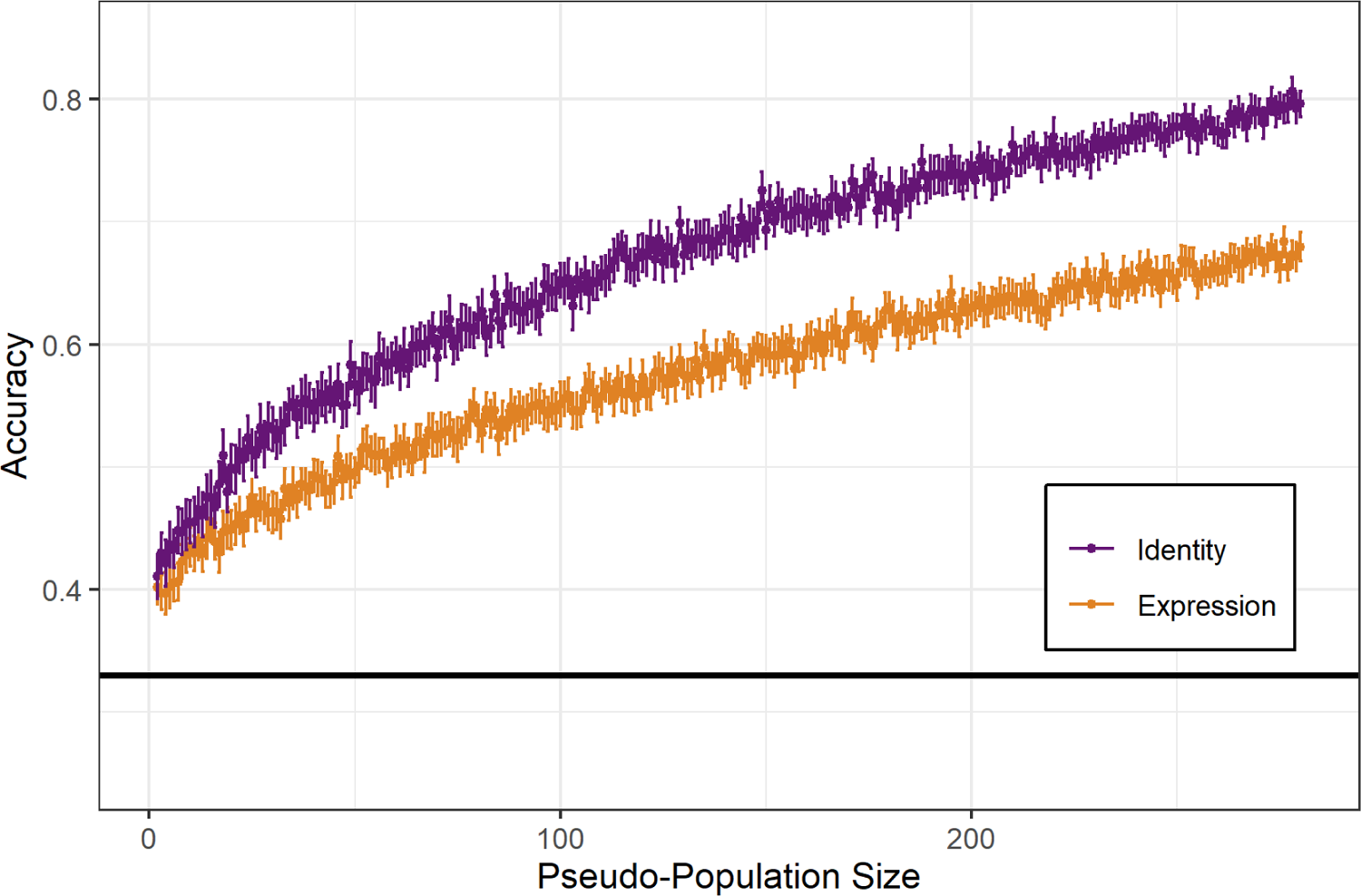
Pseudo-populations of Single Units Facilitate Better Decoding of Identity and Expression Pseudo-populations of sizes 2-280 were constructed from random samples of the 280 units. A KNN decoder was used to classify the single trials by their nearest neighbors (Euclidean distance). The maximum accuracy from classification using 5, 7, or 9 neighbors was used as the accuracy metric. At each pseudo-population size, 50 repetitions of 2-Fold CV were conducted (each repetition with a random selection of units) and averaged to produce the mean and standard error for the population size. Dots represent the mean decoding accuracy at a given population size for a particular variable, error bars represent the 95% confidence interval of the mean (± 1.96 X standard error). Decoding accuracies for both variables increase as population size increases, with identity increasing faster and ultimately having higher accuracy at larger populations than expression.

**Figure 8:**
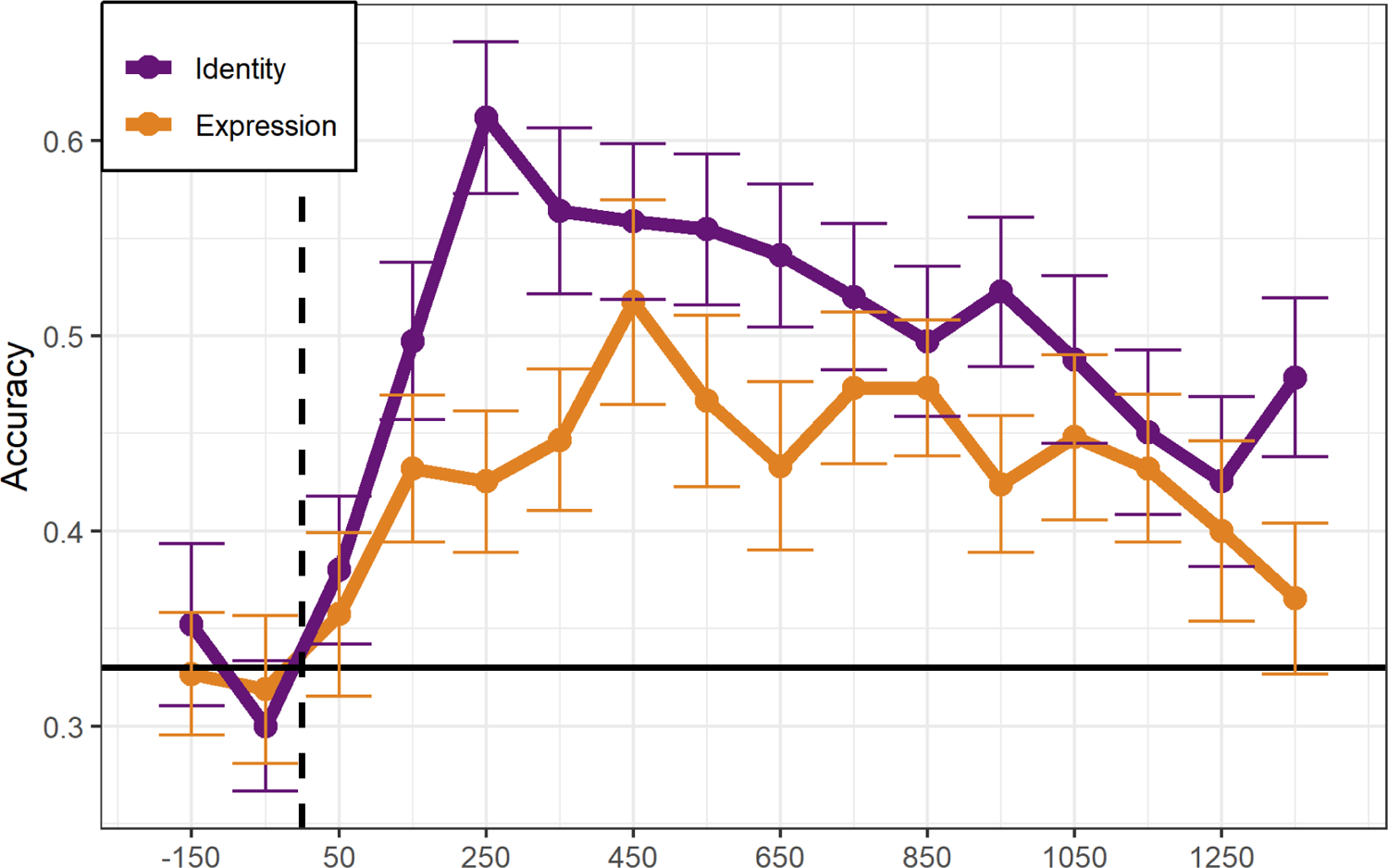
Sliding Bin Decoding Analysis Reveals Early Peak of Identity Decoding A decoding analysis was performed on 285 unit pseudo-population firing rates in 200ms bins, across the response period. Dots represent the mean decoding accuracy for the variable of interest across 50 iterations of 5-fold CV. Dots represent the center of the bin (i.e. a dot at 50ms represents a bin from −50 to 150ms). Error bars represent the 95% confidence interval of the mean (1.96 * standard error). Peak decoding of identity was higher and occurred earlier than that for expression. Additionally, the mean decoding accuracy for identity was greater than expression for all bins after stimulus onset.

Single unit responses, like our naturalistic stimuli, were dynamic across time. Considering accurate decoding could be achieved at the population level, we sought to understand the time course of information about expression type and conspecific identity across the population. To do so, we employed a time-binned decoding model, where pseudo-population firing rates were calculated for discrete, 200ms time bins across the response period. The period from 250ms before the stimulus to 1450ms after the stimulus was analyzed in 200ms bins, overlapping by 100ms. A decoding model using the maximum correlation coefficient between a single trial and mean templates of training trials was used to classify single trials based on the 285 unit pseudo-population response (5-fold CV, 50 iterations). This model was run for each 200ms bin in the stimulus period and for each variable of interest (expression and identity, see Methods: Decoding of Identity and Expression Across Time). The mean and 95% confidence interval of the decoding accuracy was produced and plotted for each time bin in Figure 8.

The mean decoding accuracy for identity was higher than that of expression for all time bins after stimulus onset, reaching a peak of 0.61 at the bin centered on 250ms post-stimulus onset. Expression decoding reached a peak of 0.51 at 450ms after stimulus onset. 95% confidence intervals for both variables were non-overlapping for bins centered on 250ms, 350ms, 550ms, 650ms, 750ms, and 1350ms. Overall, identity decoding increased earlier and to a higher level than that for expression. Decoding for both variables remained above chance for multiple bins after the peak of their decoding accuracy without a clear return to chance over our response period.

This suggests that viewing expressive stimuli generates sustained population activity in the VLPFC that continues to carry information about conspecific identity and expression type well after a stimulus has terminated. The early peak of identity decoding followed by expression decoding is consistent with the availability of this information over time in our naturalistic stimuli.

### Identity Shapes the Population Response to Naturalistic Stimuli

Across our decoding analyses, we observed earlier and persistently robust decoding of identity across the stimulus period compared to the decoding of expression (Figures 8). Given these observations, we explored the relationships of both of these categorical variables within the population activity space. In particular, we sought to understand if there was an inherent structure within this space that would facilitate accurate decoding of conspecific identity. In order to do this, we conducted a PCA on the mean firing rates of our 285 responsive units for each of the 9 stimuli (see Methods: Principal Component, Group Distance, and Trajectory Analysis). Once again, because individual units were recorded during 1 of 2 examples of an identity-expression combination, this population analysis assumes a generalization of conspecific identity and expression type across two exemplars.

The variance explained by the first 3 principal components (PCs) was 47.4% (PC1: 18.8%, PC2: 15.1%, PC3: 13.5%) without a clear elbow across the 8 PCs, suggesting the population response space is inherently high-dimensional. Nonetheless, visualization of population responses in PC1 and PC2 displayed a trend found across all 8 retained dimensions; within PCs 1 and 2, population responses to the same identity appeared to occupy non-overlapping subspaces within the response space (Figure 9A). In other words, although a population response might be closer to that of a different conspecific identity than to one of it’s own (as is the case for BK-AFL and ST-AGG), responses to the same identities could be connected together without overlapping the space occupied by other connected identities. This suggested that population responses to the same identity might be closer together than those for expression types or those for mismatching stimuli in the population space.

**Figure 9:**
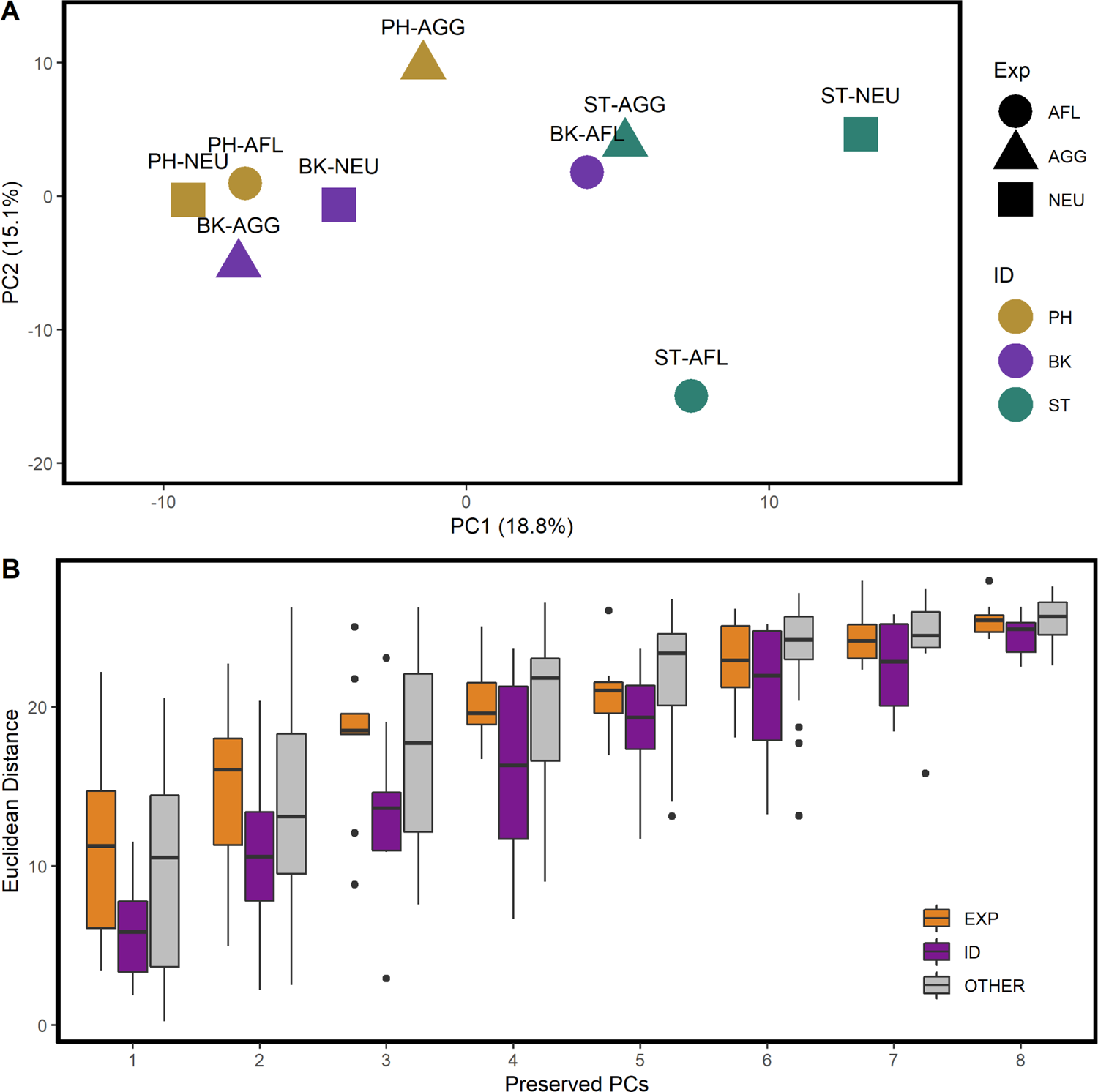
PCA Population Responses to Identity are Closer than Expressions The mean firing rates over 0-1500ms for each stimuli, for all 285 responsive units, were analyzed using PCA. In A, the population responses to each of the 9 stimuli are reconstructed in PC1 and PC2. Population responses are colored by identity, to highlight that identities appear to occupy non-overlapping portions of the overall response space. In B, the distances between populations responses to similar identities (ID), expressions (EXP), or mismatching stimuli (OTHER) were measured and grouped together in reconstructed spaces preserving PCs 1-8. There were significant effects of PCs preserved (p<0.005) and Group (p<0.005) in a Two Way ANOVA (adjusted for unbalanced groups). Tukey HSD testing showed significant differences between ID-EXP (p<0.005) and ID-OTHER (p<0.005). The difference between EXP-OTHER was not

To quantitatively characterize this phenomenon, we measured the Euclidean distances between all population responses and grouped them according whether the distance was between population responses to the same conspecific identity (ID), the same expression type (EXP), or between responses to mismatches of both expression and identity (OTHER). We conducted this analysis for population spaces ranging from PC1 to PC8 (Figure 9B). In each population space, the median distance for the ID group was lower than the median distances for EXP or OTHER. A Two-Way ANOVA model fitting distance as a function of dimensions and group was statistically significant for both factors (dimension p < 0.005, group p < 0.005, ANOVA model: distance = dimension + group, see Equation 4). Post-hoc testing with Tukey HSD revealed significant differences between ID and EXP as well as ID and OTHER (p < 0.005 for both). EXP and OTHER were not significantly different (p = 0.95). Similar trends were seen when data was analyzed for each subject independently (data not shown).

Given that population responses to the same identity tended to be closer together in the population space, we pursued an understanding of the relationships between population responses across time. We particularly sought to assess whether population responses had a consistent direction of movement towards their final positions in the population response space, based on the identity of the response. To do so, we projected the mean firing rates of the 285 neuron population response at various time points along the 0-1500ms stimulus period into a common population response space (see Methods: Principal Component, Group Distance, and Trajectory Analysis). By plotting each population response’s trajectory to its final position, we were able to visualize the average response of the population, over time, to each of the stimuli in our set (Figure 10). Viewed from multiple PC angles, a population response to a particular conspecific identity appears to emanate from a central region, common to all responses, and follow a trajectory, through time, that is similar to its identity counterparts. In views where responses to the same identity appear to splay in separate directions, orthogonal views corroborate that the appearance of splay is typically an artifact of the viewing angle (e.g. ST in Teal, Figure 10C shows splaying while A and B do not). This consistency of trajectory is not readily seen with populations responses to the same expression type (Figure 10D).

**Figure 10:**
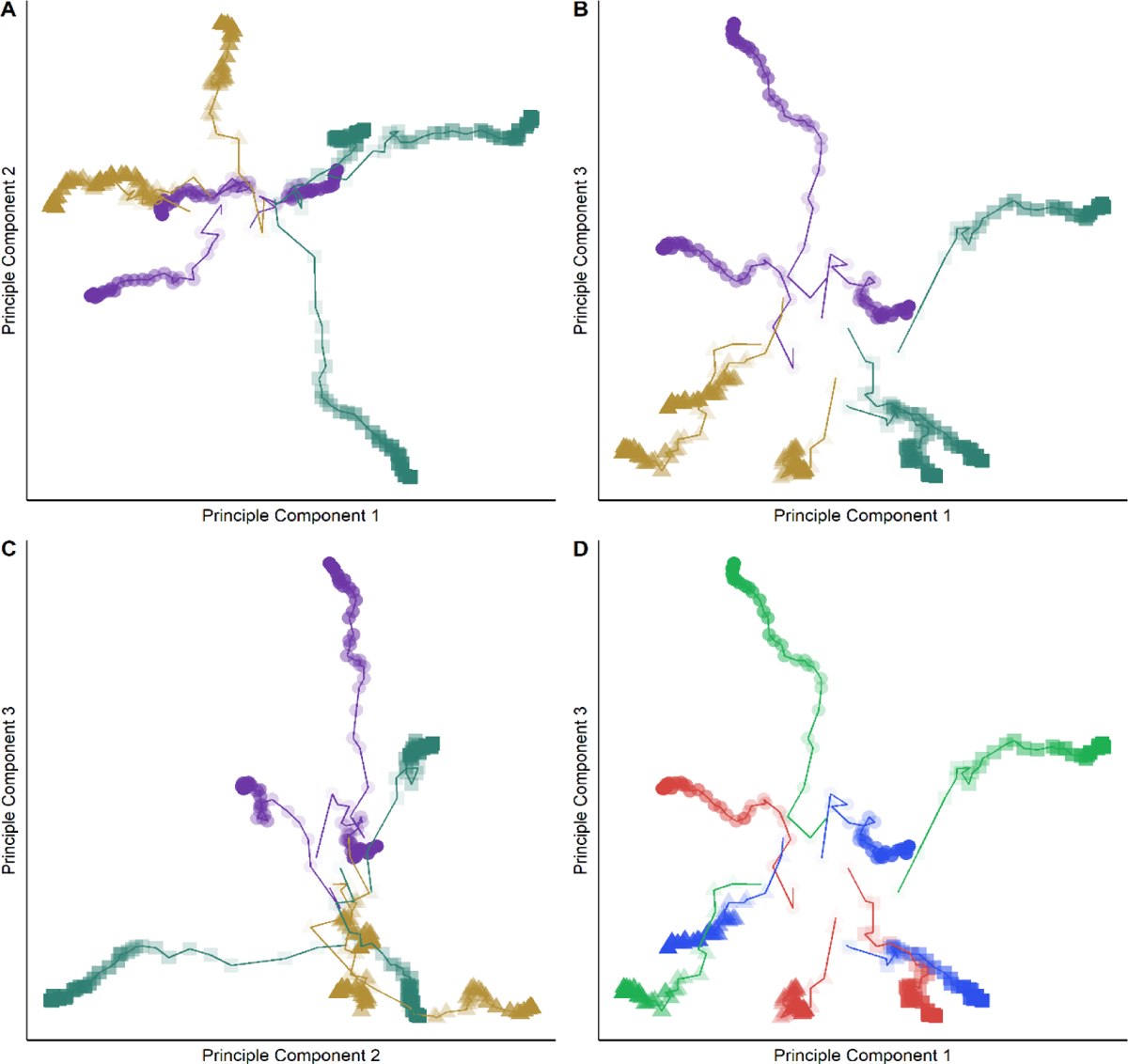
PCA Population Responses to Naturalistic Stimuli Follow Identity Based Trajectories The positions of the population response to each stimulus through time were projected onto the 0-1500ms firing rate PCA space from **Figure 11**. Coloring of identities is the same as the previous figure, markers have been adjusted to reflect the same identity as well. Each marker represents the position of the population at a point in time, for a given stimulus. The transparency of the marker is proportional to the time point, wherein very transparent markers represent firing rates across the early response period (e.g. 0-150ms) and less transparent markers represent firing rates across a longer response period (e.g. 0-1200ms). Trajectories are connected by lines for ease of following. (A) reflects a view of the population mean trajectories from PC1 and PC2. (B) is from PC1 and PC3. (C) is from PC2 and PC3. (D) is the same as (B), recolored to group expressions rather than identities (red = aggressive, blue = affiliative, green = neutral). The trajectories emanate from a central region and the trajectories of the same identity appear to move towards a common subspace.

## Discussion

Our findings indicate that, despite a diversity of responses exhibited at the single cell level, population activity in the macaque VLPFC in response to the audiovisual expressions of conspecifics is primarily organized around the identity of the conspecific attended to. Single neurons, responsive to audiovisual expression stimuli in the VLPFC, exhibited complex firing patterns that varied over time (Figure 2) and exhibited weak selectivity for specific stimuli, expression type, or conspecific identity (Figure 3). Despite low selectivity, both conspecific identity and expression type exerted a systematic influence on neuronal firing rates (Figure 4) suggesting that information about both factors is transmitted by VLPFC neurons in response to the perception of a vocalization-expression. Information from a neural population, rather than single neurons, was necessary to decode identity and expression type with appreciable accuracy (Figure 7). Specifically, the higher decoding accuracy of identity at all populations sizes as well as its earlier and higher peak response time, compared to expression type (Figure 8), support a model where identity is the primary factor determining the population activity of the VLPFC. Findings from our PCA analysis confirm that the structure of the aggregate activity from responsive neurons segregates population responses to the same conspecific identity into subspaces of the total response space (Figure 9) and that the activity of the population over the response period cohesively evolves such that responses to the same identity follow a similar trajectory across time (Figure 10). Together, these findings confirm the primacy of identity in determining the population activity of the VLPFC during perception of naturalistic expressions.

Furthermore, this activity is information rich, allowing for accurate decoding of expression type at the population level and potentially many more variables based on the diversity of responses seen at the single neuron level. These findings have direct implications for the study of circuits supporting normal social functions.

### VLPFC Population Activity Reflects the Identity of Conspecifics

Studies of the macaque VLPFC have consistently produced evidence suggesting that the VLPFC has a critical social function. Functional neuroimaging of awake macaques viewing videos of conspecifics have shown activations in the VLPFC comparable to those in the superior temporal sulcus (STS) and other regions of the temporal lobe involved in perception of expressions (Russ & Leopold, 2015; Shepherd & Freiwald, 2018). Selectively-responsive face-cells and “face patches” have been identified in the macaque temporal lobe and VLPFC Tsao et al. 2008b), which are posited to be homologous with similar patches in humans (Tsao, Moeller, et al., 2008). Furthermore, the VLPFC is also highly responsive to species-specific vocalizations presented separately (Romanski and Goldman-Rakic 2002; Romanski et al., 2005), particularly when combined with the accompanying facial gestures, as in the present experiment (Diehl & Romanski, 2014; Kuraoka et al., 2015; Romanski & Hwang, 2012; Sugihara et al., 2006). VLPFC auditory cells express high selectivity for specific calls (Romanski et al., 2005) but little responsivity to pure tones and other simple auditory stimuli (Romanski et al., 2005; Romanski & Averbeck, 2009; Romanski & Goldman-Rakic, 2002).

Our combination of single unit and population level analysis illustrates the emergence of a cohesive population structure from a variety of relatively non-selective neurons. We found virtually no neurons with a specific response for a single identity and the majority had lower selectivity for conspecific identity than for specific stimuli. Decoding of identity or expression by single units, on average, barely exceeded chance. However, integrated over time and number of neurons, the population activity of the VLPFC facilitated separation of different conspecific identities from complex, multimodal, time-varying stimuli. Importantly, we did not select only face-selective neurons in our recordings and analysis. Rather, we recorded from cells broadly across the VLPFC and assessed the activity of those that were responsive to naturalistic, audiovisual expression stimuli. As such, our results are, perhaps, more aligned with studies of the functional nature of the entire VLPFC under naturalistic conditions (Russ & Leopold, 2015; Shepherd & Freiwald, 2018) than those specifically targeting the face patches with a large variety of static face stimuli (Barat et al., 2018). Regardless, without selecting for sites or neurons specific to faces, and including the vocalizations that accompany prototypical facial gestures, our analysis showed that population level activity could be utilized to accurately decode both expression type and conspecific identity and that the population activity response followed a time evolving structure that aligned with identity. Our findings, combined with well-established anatomical connectivity between the VLPFC and temporal cortex (Barbas, 1988; Felleman & Van Essen, 1991; Kravitz et al., 2013; Romanski, Bates, et al., 1999; Romanski, Tian, et al., 1999) as well as the prevalence of identity information across the face-patch network (Chang & Tsao, 2017; Yang & Freiwald, 2021), support the idea that the VLPFC is a critical region in social communication.

### Expression Decoding Is Likely Facilitated by a High Dimensional Response Space

Despite the population level impact of conspecific identity and expression type, single neurons could not be clearly categorized as “identity” or “expression” responsive neurons due to inconsistent firing patterns across the exemplars of each category. This is consistent with studies of responses to macaque vocalizations (Romanski et al., 2005), which show that single VLPFC neurons do not encode categories across multiple exemplars. Rather, the activity patterns observed at the single neuron level resemble mixed selectivity (Rigotti et al., 2013), in which neurons have robust, typically context dependent, responses to a single or a few stimuli and are also responsive, to a lesser degree, to other stimuli in the set (Figure 5). Furthermore, VLPFC neurons exhibit non-linear multisensory integration in response to species-specific facial and vocal expressions like the audiovisual expressions used in the current study (Diehl & Romanski, 2014; Sugihara et al., 2006), which fundamentally increases the dimensionality of the population response space beyond those of the two sensory modalities (Rigotti et al., 2013). Thus, it is highly likely that the idiosyncratic response patterns seen at the single unit level are a result of mixed selectivity of sensory information that is non-linearly integrated by VLPFC. The purported utility of non-linear mixed selectivity is the simultaneous existence of a very high dimensional population response space capable of encoding a variety of unique stimuli while also providing redundant and reliable encoding that forgoes the need for neurons that are specifically responsive to a particular stimulus feature of interest (Johnston et al., 2020; Rigotti et al., 2013).

Our ability to decode expression above chance is likely the result of the existence of a high dimensional space created by non-linear integration in the VLPFC. Even though we find that conspecific identity is the dominant variable structuring the population response in the first three principal components, we were able to decode expression with increasing accuracy as more neurons were added to the population (Figure 7). Thus, while population responses to expression types seem to be diverging in lower dimensions, our decoding model (which relies on proximity in space) suggests that expression may be converging in higher dimensions.

An additional factor facilitating the decoding of expression is our use of naturalistic stimuli with synchronous, dynamic auditory and visual information that affords additional features in the time domain. For example, the expansion and contraction of the mouth presented with a simultaneous rapid increase in pitch of a vocalization may be a feature encoded by these neurons. Most neurophysiology studies of expression and identity in the macaque employ static visual stimuli (Barat et al., 2018; Gothard et al., 2007; Tsao, Schweers, et al., 2008), present an auditory stimulus with a static image (Barat et al., 2018), and/or separate the visual movie from the vocalization (Kuraoka et al., 2015; Kuraoka & Nakamura, 2007). These paradigms afford greater control of stimulus timing and increase the variety of stimuli at the expense of failing to recapitulate naturalistic multisensory integration and dynamic multisensory features. Our methods are more aligned with fMRI studies of the macaque VLPFC, in which viewing of naturalistic videos (without associated audio) revealed that motion is a critical correlate of activation and that the activation is specific to social-motion (Russ & Leopold, 2015).

As a result of our approach, our stimuli were constrained by the natural duration of the audiovisual expression and by what exemplars could be elicited from live subjects, such that subtle features (direction of gaze or the amplitude of the mouth movement) could not be controlled for without degrading the natural quality or timing of the expressions. Fortunately, recent studies of macaque social perception (Khandhadia et al., 2021) have started employing natural audiovisual stimuli and naturalistic 3D macaque avatars for which specific features of expression, gaze, and facial action can be parametrically manipulated within a single identity (Murphy & Leopold, 2019; Wilson et al., 2020). Further single unit and population-level studies employing these methods are needed to understand the specific correlates of expression that are encoded in the activity of neurons in the social communication network.

## Acknowledgements

We would like to thank Oishee Raman and John Housel for their technical assistance and expertise in spike-sorting, animal training, and histology. We would also like to thank Marc Scheiber for his guidance and assistance with the FMA surgical procedures. This study was supported by NIH DC 04845 (LMR), The Schmitt Program for Integrative Neuroscience (LMR), F30 MH122048 (KKS), and the University of Rochester Medical Scientist Training Program (T32 GM007356).

## Notes

**Conflict of Interest Statement:** All authors have no conflicts of interests to disclose.

### Competing Interest Statement

The authors have declared no competing interest.

### Summary of Updates

Combination of Figures 2 and 3, revision of manuscript to fit Journal of Neuroscience word limits, removal of supplemental data and references to data not integral to the body of work presented.

